# Plasticity of *Medicago truncatula* seed dormancy relates to large-scale environment variation

**DOI:** 10.1101/2019.12.22.886218

**Authors:** Juan Pablo Renzi, Martin Duchoslav, Jan Brus, Iveta Hradilová, Vilém Pechanec, Tadeáš Václavek, Jitka Machalová, Karel Hron, Jerome Verdier, Petr Smýkal

## Abstract

Seed dormancy and timing of its release is important developmental transition determining the survival of individual as well as population and species. We used *Medicago truncatula* as model to study legume seed dormancy in ecological and genomics context. The effect of oscillating temperatures as one of the dormancy release factor was tested over the period of 88 days on the set of 178 accessions originating from variable environmental conditions of Mediterranean basin. Phenotypic plasticity of final dormancy was significantly correlated with increased aridity, suggesting that plastic responses to external stimuli provide seeds with strong bet-hedging capacity and the potential to cope with high levels of environmental heterogeneity. Genome-wide association analysis identified candidate genes associated with dormancy release related to secondary metabolites synthesis, hormone regulation and modification of the cell wall likely mediating seed coat permeability and ultimately imbibition and germination.

**Highlight:** Medicago seed dormancy was correlated with increased aridity of the environment, suggesting that plastic responses provide seeds with a bet-hedging capacity. Genome-wide association analysis identified candidate genes associated with release from dormancy.

## Introduction

Plant species exhibit a high level of local adaptation and phenotypic plasticity may contribute to the distribution range. While the local adaptation is the genetically fixed advantage of a population under certain environmental conditions (Tognetti *et al*., 2019), the phenotypic plasticity is the ability of a genotype to generate different phenotypes according to the environment (Valladares *et al*., 2006). This variation is created by interplay of genetic and environmental factors. Understanding of the genetic basis of local adaptation and phenotype plasticity is relevant for climate change, crop production, conservation and understanding of the speciation. Adaptation to local environment is a primary force driving evolution, as demonstrated in transplantation and contemporary studies (Clausen, 1951; Levin, 2000). On the other hand, phenotypic plasticity may allow species to growth and survive in different environments despite a restricted genetic base, and thus could be advantageous under highly variable environments including climatic change (Nicotra *et al*., 2010). In plasticity studies, trait changes under different environmental conditions and this response is quantified (Valladares *et al*., 2006). The complimentary evaluation between phenotypic plasticity and local adaptation can be performed by the norms of reactions. The slope of the trait value across a set of environments indicates the level of plasticity of each genotype, and significant genotypes × environment interaction suggests local adaptation (Tognetti *et al*., 2019).

The development of remote sensing technologies and available environmental data derived from geographic information systems (GIS) provide information on the patterns of environmental variation at continental scale. These data are used to identify environmental differences between populations and species (Kozak *et al*., 2008), and to infer ecological and evolutionary processes from these patterns (Bragg *et al*., 2015). Examples of application of association between genome-wide and eco-geographical data include salinity (Friesen *et al*., 2014), rhizobium symbiosis (Stanton-Geddes *et al*., 2010), soil (Guerrero *et al*., 2019) and identification of genomic signature of *M. truncatula* (Yoder *et al*., 2014) or *Arabidopsis thaliana* (Hancock *et al*., 2011) adaptation to the climate.

Timing of the seed germination is one of the key steps in plant life, influencing subsequently destiny of individual as well as whole population at the given place. Plants have evolved various mechanisms to control the entry of the quiescent seed protecting embryo into vulnerable environment (Baskin and Baskin, 2014). These controls are finer in those annual species that do not frequently and reliably reproduce, because seed dormancy is a key trait enabling them to persist in an unpredictable environment (Duncan *et al*., 2019).

Beside genetic factors, seed germination is regulated by environmental factors (Finch-Savage and Leubner-Metzger, 2006; Graeber *et al*., 2012; Nonogaki, 2014). Seasonal and diurnal fluctuations in temperature are considered the main regulatory factors of seed dormancy (Bewley *et al*., 2013; Probert 2000), while in arid regions with high inter-annual rainfall variability, soil moisture is a major determinant for seedling emergence (Holst *et al*., 2007).

A diverse range of dormancy mechanisms (blocks) has evolved in relation with the diversity of climates and habitats (Willis *et al*., 2014). Three main dormancy classes have been proposed (Baskin and Baskin, 2014; Willis *et al*., 2014). 1) Morphological dormancy (MD) refers to seeds that have an underdeveloped embryo and require time to grow. 2) Physiological dormancy (PD), involves abscisic acid and gibberellins metabolism. 3) Physical dormancy (PY) is caused by water-impermeable palisade cells in seed (or fruit) coats. This type of dormancy occurs in at least 18 angiosperm plant families and is particularly common in legumes (Hradilová *et al*., 2019; Norman *et al*., 2002; Smýkal *et al*., 2014). It is unknown how and why PY varies inter- and intra-specifically in natural ecosystems (Hudson *et al*., 2015).

The prevention of germination of certain proportion of seeds reduce the risk of extinction once conditions turned to unfavourable, especially under less predictable environmental conditions. The study of Rubio de Casas *et al*. (2017) showed that legume seed dormancy is linked with environment seasonality. It has been suggested theoretically (Templeton and Levin, 1979) and shown empirically (Evans *et al*., 2007) that adaptation for dormancy is a bet-hedging strategy to magnify the evolutionary effect of “good” years and to dampen the effect of “bad” years, i.e. to buffer environmental variability (Venable, 2007). In addition, species that frequently and reliably produce seed can afford riskier germination under unfavourable conditions (e.g. small rainfall events) because the consequences of failure to establish are less dire than for species that do not reliably produce seed (Duncan *et al*., 2019).

There are elaborate checks on germination requiring particular environmental conditions to break seed dormancy and elicit germination (Baskin and Baskin, 2014). However, little is known about clues which release the PY dormancy. By experimental testing, beside scarification, wet or dry heat were found effective (Baskin and Baskin, 2014; Hradilová *et al*., 2019; van Klinken *et al*., 2006). It is hypothesized that and in natural conditions the temperature and soil moisture oscillations are also the major players (Batlla and Benech-Arnold, 2015; Ellis *et al*., 1987). Laboratory studies have demonstrated an association between seed responsiveness to temperature and the thermic characteristics of their habitat range (Hradilová *et al*., 2019; Renzi *et al*., 2018; Rosbakh and Poschlod, 2015).

*Medicago truncatula* (barrel medic) is an annual, diploid, self-fertile and prolific seed production species with a natural geographic distribution across the Mediterranean Basin. Phenotypic variation among populations is potentially explained by adaptation to local environmental conditions (Ronfort *et al*., 2006). It offers excellent model to study seed dormancy in relation to genetic and environmental factors. *Medicago truncatula* seeds exhibit both physical and physiological dormancy. Physiological dormancy (PD) in *M. truncatula* seeds is non deep, and is removed during seed ripening period (Gallardo *et al*., 2006). The short after-ripening period to overcome PD (< 3 month) determines that PY release is the most important trait to regulate the timing of seedling emergence. Despite this, most germination studies in *M. truncatula* eliminate the influence of PY dormancy through prolonged periods of stored (> 9 months) and/or by seed scarification (Faria *et al*., 2005; Brunel *et al*., 2009; Vu *et al*., 2015).

Large georeferenced collections (Ronfort *et al*., 2006), reference genome and a high-density SNP map of more than 260 genotypes (Stanton-Geddes *et al*., 2013) are available for this species and was used to study association between genome and environment in relation to flowering (Burgarella *et al*., 2016; Julier *et al*., 2007; Stanton-Geddes *et al*., 2013).

The present study aimed to analyze pattern of dormancy release of 178 accessions of *Medicago truncatula* originating from various environments of Mediterranean basin to oscillating temperature treatments. Following main questions were addressed: 1) Is there variation in physical seed dormancy among accessions to temperature treatments? 2) Which of the ecological factors acting as potential adaptation drivers are correlated with dormancy? 3) Is there association between candidate loci and level of *Medicago* seed dormancy using genome-wide analysis?

## Material and Methods

### Plant material

Seeds of *Medicago truncatula* accessions were retrieved from INRA Montpellier (http://www1.montpellier.inra.fr) and from University of Minnesota, USA (http://www.medicagohapmap.org/hapmap/germplasm). These accessions were selected from HapMap collection (Burgarella *et al*., 2016; Yoder *et al*., 2014) with subtracted accessions with insufficiently accurate GPS data. Plants were grown in glasshouse conditions at Department of Botany, Palacký University, CZ from March to July (2016) and from September to May (2017, 2018). Plants were cultivated in 3 litre pots with sand peat substrate (1:9) mixture (Florcom Profi, BB Com Ltd., CZ), watered as required and weekly fertilized (Kristalon Plod a Květ, Agro, CZ). Temperature varied according to weather from minimum of 15°C during winter time to maximum of 40°C in late spring. Supplementary light was provided (Sylvania Grolux 600 W, Hortilux Schreder, Holland) to extend the photoperiod to 8h during September-February and to 14h from February on to stimulate the flowering. The mature pods were collected, packed in paper bags and dried at 20°C and 60–63% relative humidity to allow post ripening for period of 4 to 6 weeks prior testing. Sufficient seed stock was obtained from 178 accessions using in house made equipment (https://www.youtube.com/watch?v=y0_4h9LYEhA).

### Seed dormancy release experiments

Release of seed dormancy was tested as imbibition (e.g. uptake of water) and terminated when radicle protruded seed coat (Hradilová *et al*., 2019). As our study was aimed to study PY, the values we used in all subsequent analysis correspond to imbibed seeds. To mimic natural conditions, two temperature treatments (oscillating temperatures of 35/15°C and 25/15°C in the dark at 14h/10h (day/night)) were applied to intact seed batches (50 seeds, in 2 to 3 replicas per treatment). Seeds were placed onto water saturated filter papers (Whatman Grade 1, Sigma, CZ) in 60 mm Petri dishes (P-Lab, CZ) in temperature controlled chambers (Laboratory Incubator ST4, BioTech, CZ). In order to prevent fungal growth (as seeds sterilization would alter seed coat properties) fungicide (Maxim XL 035 FS; containing metalaxyl 10 g and fludioxonil 25 g) was applied. Seeds were monitored at 24h intervals for total of 88 days. After each scoring, the plates were randomly relocated within chambers. Germinated seeds (e.g. when radicle protruded from testa) were removed. At the end of the testing, remaining seeds were scarified and let to germinate to verify their viability. This typically was over 98%. Although we have selected macroscopically intact seeds for experiments, we cannot exclude some microscopic cracks on seeds as results of mechanical damage during the threshing procedure. We observed that in the course of first hours of seed germination experiments, certain proportion of the seeds imbibe. Therefore, we have subtracted the first day imbibition value from the analysis detailed bellow. In 2016, 2017 and 2018 year 147, 74 and 130 accessions were tested (Figure S1). In total, seeds of 178 accessions were included in the experiments (Table S1).

### Evaluation of dormancy and germination traits

Original germination data (daily counts of imbibed seeds representing a finite sample from theoretically infinite population) were considered as discrete realizations of an asymptotically continuous process, approximated by spline functions (de Boor, 1978; Machalová, 2002). The resulting smoothing spline called the *absolute germination distribution function* (AGDF; as applied in pea by Hradilová *et al*., 2019) was used in analysis. Accordingly, the area under curve (AUC) of the spline function takes into account both the course of the germination as well as the final score of germinated seeds which capture the dynamics of seed germination better than existing germination coefficients (Kader, 2005; Talská *et al*., submitted).

Several statistics (traits) characterizing dynamics and final state of dormancy release of seeds of each accession for each treatment (i.e. 25/15°C and 35/15°C) were calculated. (i) Final PY dormancy (%; FPYD_25_, FPYD_35_): represents percentage of dormant seeds at the end of experiment after excluding seeds germinating during the first day of each experiment (i.e., 100 – final germination percentage after 88 days + germination percentage after first day), calculated separately for two germination treatments. (ii) Germination pattern (AUC_25_, AUC_35_): this trait represents the area under curve (AUC) coefficient that takes into calculation both dynamics of germination as well as final germination percentage (see above). High values mean rapid and early germination of majority of seeds. (iii) FPYD_M_ and AUC_M_: these two coefficients represent means of respective coefficients estimated separately for each temperature treatments (e.g., FDYD = (FPYD_25_+FPYD_35_)/2)), (iv) Germination response (AUC_35-25_): this is the difference between two AUC (i.e., AUC_35_-AUC_25_) of the same accession calculated for two germination treatments (i.e. 35/15°C and 25/15°C). Higher absolute value of germination response means larger difference in germination pattern of the same accession between germination treatments while sign of the difference suggests which of the treatment shows larger AUC. (v) Phenotypic plasticity index of the germination pattern (PI_AUC_) and (vi) Phenotypic plasticity index of final PY dormancy (PI_PY_): these traits were calculated for each accession as (trait_max_−trait_min_)/trait_max_, where trait_max_ and trait_min_ were respectively the maximal and minimal value of the trait measured on the same accession across the two temperature treatments (25/15°C and 35/15°C). This estimate characterize the maximal plastic capacity of an individual in variable environments without taking into account the direction of the plastic response or the change in intensity with environmental variation. Phenotypic plasticity index range from 0 (no plasticity) to 1 (maximal plasticity) (Valladares *et al*., 2006).

For the purpose of multivariate analyses, we calculated average of each dormancy trait for each accession over the years. Consequently, the matrix of averages of each dormancy trait for each accession was used in multivariate analyses. Multicollinearity among variables was assessed by variance inflation factor (VIF) for quantitative traits using library usdm (Naimi *et al*., 2014) in R. Only variables whose VIF was lower than 15, were retained in the analyses. Except for FDYD_M_ and AUC_M_, any of abovementioned traits had collinearity problem.

### Environmental variables and spatial accuracy

Due to different spatial accuracy of accessions and in order to minimize the spatial error caused by imprecise coordinates we developed a geoprocessing model in ArcGIS PRO environment (https://pro.arcgis.com/). Model automated the calculation of mean values of selected variable from 5 km buffer around the each collection site in order to smooth the uncertainty caused by imprecise localization.

#### Climatic variables

The WorldClim database (http://worldclim.org) version 2.0 was used to extract climatic data (period 1970-2000) from GeoTIFF rasters in the WGS-84 coordinate system (EPSG: 4326) with a spatial resolution of 30 arc-seconds (∼ 1 km). Bioclimatic variables (BIO1–BIO19) are derived from the monthly temperature and rainfall values (Fick and Hijmans, 2017), and represent annual trends (e.g., mean annual temperature BIO1, annual precipitation BIO12), seasonality (e.g. annual range in temperature and precipitation BIO4 and BIO15) and extreme or limiting environmental factors (e.g., the temperature of the coldest and warmest month BIO5 and BIO6, and amount of precipitation in the wet and dry quarters BIO16 and BIO17).

In order to determine the inter-annual variability in selected bioclimatic variables (BIO1, BIO5, BIO10, and BIO12) between the period 1981-2010 the index of variability (*IV*) was calculated following the percentile-analysis method (Cobon *et al*., 2019). To obtain yearly mean values for years 1981–2010, we used 2[m air temperature (Kelvin degrees) and 2[m specific humidity (kg of water/kg of air) hourly data from the Modern Era Retrospective Analysis for Research and Applications Reanalysis (MERRA) 2D Incremental Analysis Update atmospheric single-level diagnostics product (MAT1NXSLV), provided by the NASA Global Modelling and Assimilation Office (Vega *et al*., 2018). Data were interpolated in spatial resolution of 2.5 arc-minutes. Temperature data were converted to degrees of Celsius (BIO1, 5 and 10). Resulting BIO12 (1981–2010) describes the annual mean of specific humidity instead of cumulative annual rainfall. Twelve month means (BIO1, 5, 10 and 12) for each site was calculated as follow: *IV* = [(90^th^ percentile - 10^th^ percentile)/50^th^ percentile]*10

The different *IV* classes vary from low (*IV* <0.50), low–moderate (0.50–0.75), moderate (0.75–1.00), moderate–high (1.00–1.25), high (1.25–1.50), very high (1.50–2.00), and extreme (*IV* >2.00).

#### Soil data

Soil data were extracted from SoilGrids collection of soil property and class maps of the world (Hengl *et al*., 2017). The predictions are based on globally fitted models of soil profile and environmental covariate data at 1 km / 250 m spatial resolutions using automated soil mapping based on machine learning algorithms.

### Testing of relationships among dormancy traits, geography and environmental variables

Matrix of environmental variables was checked for the presence of the multicollinearity using VIF. The reduced set of environmental variables (with VIF < 15), including 14 bioclimatic variables and 8 soil variables, was used in all further analyses. For each pair of variables, bivariate scatter plots together with fitted loess smooth line were displayed and Pearson’s Correlation Coefficient was calculated using library PerformanceAnalytics (Peterson *et al*., 2019) in R.

The matrix of reduced-set of environmental variables was analysed by Principal Component Analysis (PCA; Legendre and Legendre, 2012) using Canoco 5.10 (ter Braak and Šmilauer, 2012) to find the main environmental gradients within the dataset. Several precipitation variables were log(x+1) transformed and subsequently each variable was standardized to zero mean and unit variance before PCA. Set of germination traits and geographic coordinates (latitude, longitude) were used as supplementary variables and correlated with the first two principal components. To control for possible spatial autocorrelation between each germination trait and principal components representing environmental gradient, modified version of the t-test (Dutilleul *et al*., 1993) performed in SAM 4.0 (Rangel *et al*., 2010) was used. To assess whether there is spatial autocorrelation present in the PCA scores along first two axes and dormancy traits, Moran’s I spatial correlation statistics (Legendre and Legendre, 2012) was calculated for each variable using PASSaGE v. 2.0 (Rosenberg and Anderson, 2011). Ten distance classes with equal widths were created and Moran’s I and its 95%CI were calculated for each distance class.

*Medicago* accessions were grouped by macro-environmental clusters (Table 1S) based on Euclidean distance of environmental variables. Agglomeration was performed using Ward’s minimum-variance linkage algorithm. Before clustering, all variables were standardized to zero mean and unit variance.

### Phenotypic plasticity by macro-environmental clusters

We used accessions that had been repeatedly tested in 3 years (Table 1S) to calculate a norm of reaction for final PY dormancy (FPYD_25 and 35_). Final PY dormancy was used as it was the only trait that showed a significant association with the environment conditions (see results). The quantification of a norm of reaction is conceptually quite simple, where each line represents the data for a different cluster and the effect of ‘environment‘ (treatments, 25/15°C and 35/15°C). The difference between Pl_PY_ and reaction norm is expressed as the slope of the response and direction (Valladares *et al*., 2006; Sadras *et al*., 2009). To focus on the change trait in response to temperature we analysed the FPYD differences between clusters per each year by ANOVA using the InfoStat software (Di Rienzo *et al*., 2013).

### Genome-wide association analysis

Genome-wide association analysis was performed on seven seed dormancy traits (PY_25, PY_35, AUC_25, AUC_35, DEF35-25, PLAS_PY, PLAS_AUC) and three bioclimatic variables (BIO1, BIO9, BIO12) on 178 accessions. Prior to GWA analyses, normal distribution of each trait was checked using Shapiro-Wilk test. Two contrasted algorithms were used to test markers-traits associations: EMMA, a classical mixed linear model (MLM) for single locus GWAS (Yu *et al*., 2006) and FarmCPU, a multi-locus method combining the fixed effect model and random effect model iteratively in order to improve statistical power of MLM methods (Liu *et al*., 2016). Both algorithms were implemented in the R package (https://github.com/XiaoleiLiuBio/rMVP) using default parameters (p-value threshold 0.01) and run using a SNP dataset containing 5.85 million SNPs remapped in the *Medicago* genome version 5 (Pecrix *et al*., 2018). Population structure, calculated using STRUCTURE by Bonhomme *et al*. (2014) was used as covariable. Normal distribution, QQ plots and single/multiple Manhattan plots were performed using R package rMVP (https://github.com/XiaoleiLiuBio/rMVP). *Medicago* genome version 5.0 of A17 genotype (https://medicago.toulouse.inra.fr/MtrunA17r5.0-ANR) was used to search for the encoded genes within the region of 10 kb from detected SNP. Transposable elements were excluded from the search. To link identified QTNs with putative causal gene by considering the linkage disequilibrium (LD), we selected all SNPs correlated (r^2^>0.7) with top identified QTNs within a 15kb genomic range, corresponding to the average LD block size present in the Medicago Hapmap population (Bonhomme *et al*., 2013; Branca *et al*., 2011) and we listed gene IDs closely related to these SNPs. Seed expression pattern of the candidate genes was assessed using published *Medicago* seeds or seed coat expression studies (Fu *et al*., 2017; Verdier *et al*., 2013) and web based expression atlas (http://bar.utoronto.ca/efpmedicago/cgi-bin/efpWeb.cgi).

## Results

### Responses of dormancy traits of *Medicago* accessions to experimental temperature treatments

To evaluate germination and PY-dormancy, we considered seven traits in set of 178 *M. truncatula* accessions. In 2016, 2017 and 2018 year 147, 74 and 130 accessions were tested (Fig. S1, Table S1). Forty seven accessions were tested in all three years, while 129 accession in at least two years. These were tested at oscillating 35/15°C and 25/15°C temperature regimes over the period of 88 days.

Most dormancy traits exhibited a near normal distribution usually (slightly) skewed and a wide range of variability (Fig.1, Fig. S2). FPYD ranged from 34 to 100% with mean (±SD) 80±15% at 25/15°C treatment, and from 4 to 94% with mean 60±19% at 35/15°C treatment. Comparison of responses of each accession to two temperature treatments showed remarkable effect of larger temperature oscillation on dormancy release in majority of accessions (Fig. 2). The germination pattern (AUC_M_) ranged from 3 to 79, and, similarly to FPYD, larger temperature oscillation increased the dormancy release (AUC_35_: 34±17, range 5-82; AUC_25_: 24±13, range 0-79), except for some accessions (16%) where the differential (AUC_35-25_) was negative (Fig. 1, S2). Both phenotypic plasticity indexes showed large ranges with mean PI_AUC_ being slightly higher (0.56±0.29, range 0.00-1.00) than mean PI_PY_ (0.43±0.23, range 0.00-0.91) (Fig. S2).

**Figure 1.**
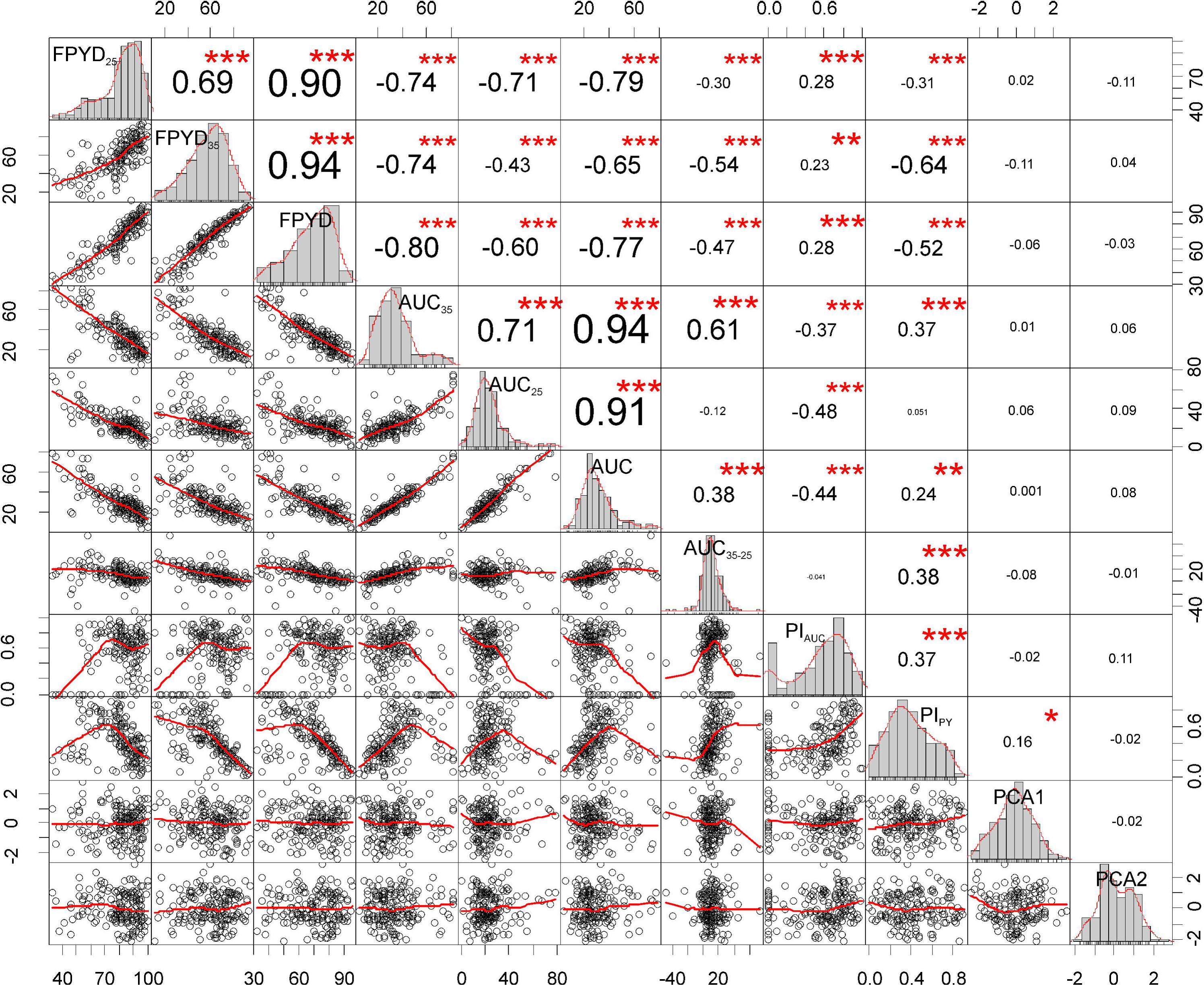
Correlation chart of dormancy release traits and ordination scores of environmental PCA of all accessions (first two ordination axes; PCA1, PCA2). The distribution of each variable is shown on the diagonal. On the bottom of the diagonal the bivariate scatter plots with a fitted loess smooth line are displayed. On the top of the diagonal the value of the Pearson correlation coefficient plus the significance level as stars are displayed. Variables BIO14, 18 and 19 were log(x+1) transformed before analyses.

**Figure 2.**
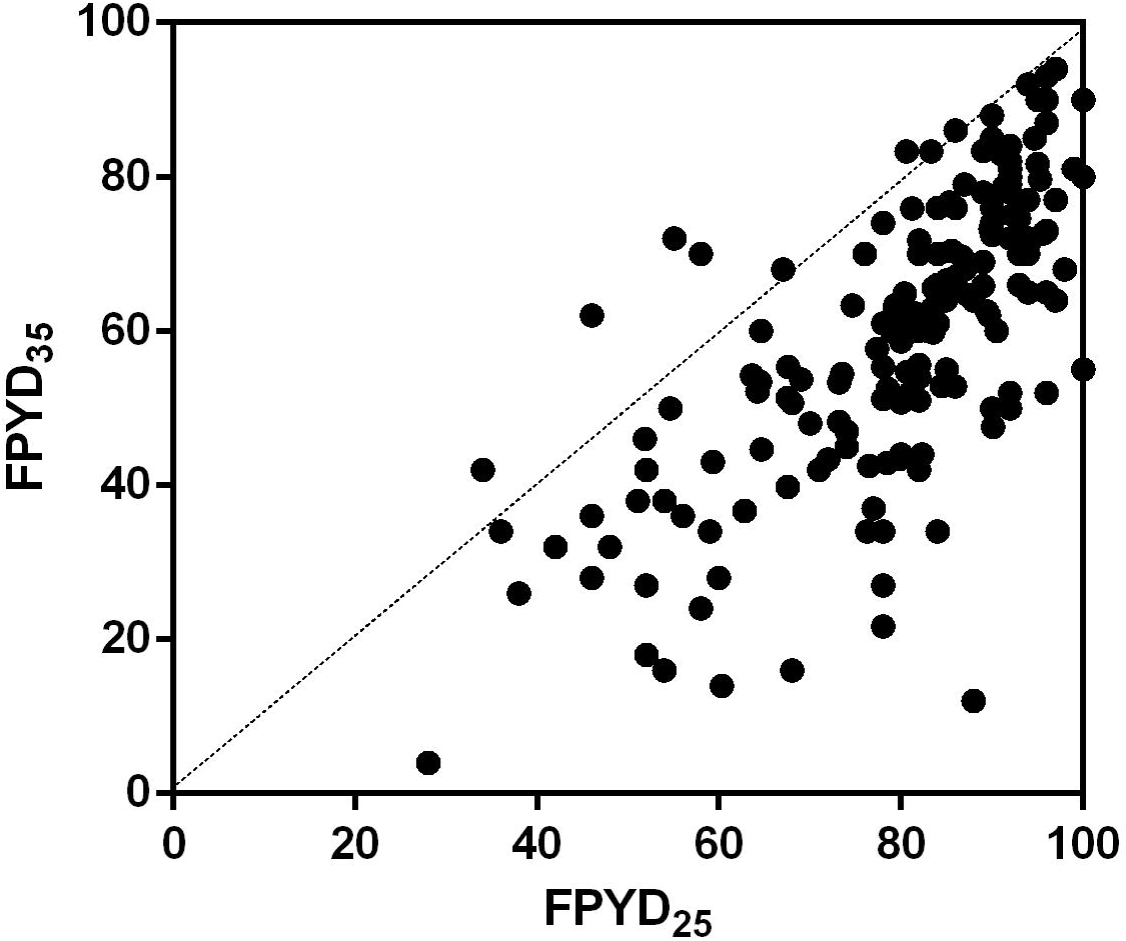
Relationship between final PY dormancy of each accession under two temperature treatments (FPYD_35_ and FPYD_25_).

To determine the relationships among different dormancy traits, we carried out correlation analyses (Fig. 1, S3, S4). All dormancy traits were moderately to strongly correlated (up to |0.75|, excl. FPYD_M_ and AUC_M_ with some correlation up to |0.94|), except for PI_AUC_ that was significantly correlated only with AUC_35_ and AUC_25_.

### Associations of environmental gradients with dormancy traits of *Medicago* accessions

Set of accessions originate from rather contrasting climatic conditions (Table S2). Mean annual temperature ranges from ca 9 to 22 °C and annual precipitation ranges from 154 to 1028 mm; consequently, temperature annual range (min-max) is from 13 to 35°C. Some accessions originate from sites with minimal winter temperatures below zero while maximal temperature of warmest months is rather similar among accessions. However, sites differ considerably in precipitations of driest and warmest periods, with many sites having zero. Concerning soil variables, considerable variability among sites was found in volumetric percentage of coarse fragments (CRFVOL) and soil organic carbon content (ORCDRC) while other soil variables were less variable. The basic descriptive statistics of the index of variability (*IV*) for BIO1, BIO5, BIO10 and BIO 12 are present in the Table 2S. *IV*BIO1 ranged from 0.33 to 1.60 with mean (±SD) 0.77±0.20, *IV*BIO5 from 0.57 to 2.26 with mean (±SD) 1.01±0.32, *IV*BIO10 from 0.63 to 1.76 with mean (±SD) 0.98±0.20, and *IV*BIO12 from 0.46 to 2.03 with mean (±SD) 0.96±0.28. The most variable *IV* was *IV*BIO5 (range 1.69) followed by *IV*BIO12 (range 1.57).

PCA of reduced data set containing 14 climatic and eight soil variables revealed two clear environmental gradients (Fig. 3). First ordination axis explained 30.4 % of the total variation and can be interpreted as gradient of aridity that is tightly correlated with latitude (i.e. north-south gradient). Climatic variables with the highest positive/negative correlation with the fist ordination axis represent temperatures of warmest month (BIO5; r= 0.61***) and driest quarter (BIO9, r = 0.71***), isothermality (BIO3, r = 0.59***), precipitation of driest month (BIO14; r = -0.83***) and precipitation of warmest quarter (BIO18, r = -0.78***). Concerning soil variables, pH index is positively correlated (r =-0.69***) while soil organic carbon content (ORCDRC, r = -0.75***), available soil water capacity (AWCh1, r = - 0.75***) and saturated water content (AWCtS, r=-0.79***) are negatively correlated with the first axis. Latitude (r= -0.77***) but not longitude (r = 0.07) is strongly negatively correlated with the first axis. Second ordination axis explained 17.3% of the total variation and can be interpreted as gradient of seasonality, with weak geographic (i.e. west-east) trend (latitude: r = -0.01, longitude: r = 0.24***). Three most correlated variables contain temperature seasonality (BIO4, r= -0.70***), precipitation seasonality (BIO15, r = 0.61***) and minimal temperature of coldest month (BIO6, r= 0.63***). Both synthetic environmental variables (PC1, PC2) are spatially structured as revealed by Moran’s I correlogram (both P << 0.001), showing positive autocorrelation at short and large distance classes and negative autocorrelation at intermediate distance classes (Fig. 3).

**Figure 3.**
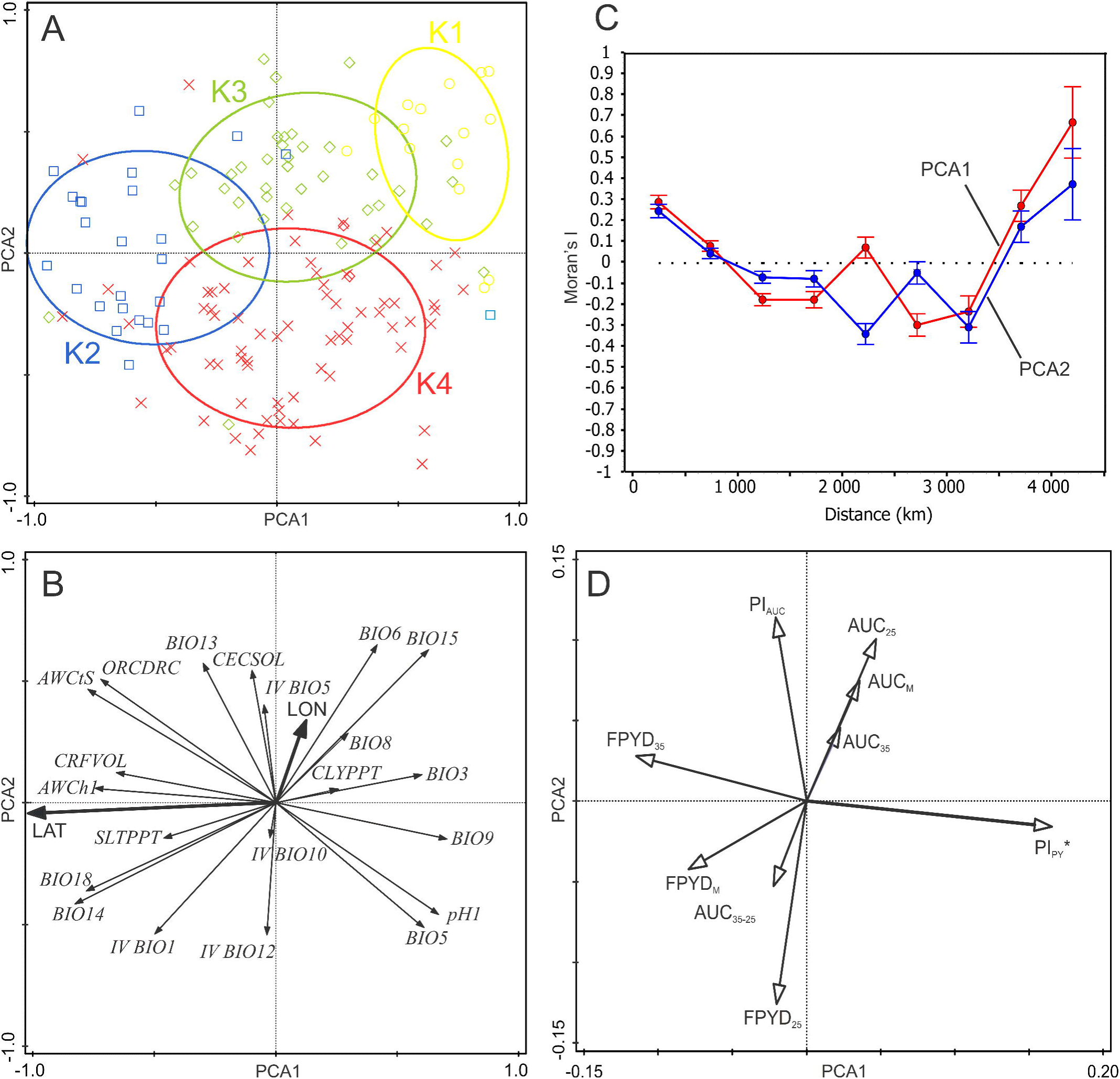
Principal component analysis (PCA) of selected bioclimatic and soil variables of *Medicago* accessions and multiple correlations of dormancy traits with ordination axes. A, B) Principal Component analysis (PCA) of selected bioclimatic and soil variables of *Medicago* accessions. Each accession is classified according to cluster analysis of environmental variables into one of four clusters (see text). The ellipses were created based on a model of bivariate normal distribution of the cluster class symbols (estimated from a variance-covariance matrix of their X and Y coordinates) to cover 95% of that distribution’ cases. Vectors of geographic variables (latitude, longitude) are added after PCA to visualise spatial gradients of environment. C) Spatial autocorrelation diagram of Moran’s I for the first two ordination axes of PCA (PCA1, PCA2). Mean ± 95%CI of I for respective distance class is calculated. D) Multiple correlations of dormancy traits with the first and the second ordination axes of the PCA. Each arrow points in the direction of the steepest increase of the values for corresponding dormancy trait. The angle between arrows indicates the sign of the correlation between the variables. The length of the variable arrows is the multiple correlation of that variable with the ordination axes. Dormancy trait significantly correlated (P ≤ 0.05, spatial correlation) with any ordination axis has an asterisk.

Inspection of dormancy traits correlation with ordination axes representing synthetic environmental variables showed that only PI_PY_ was significantly correlated with the first ordination axis (r = 0.16*), even after correction for spatial autocorrelation (P = 0.032). Other dormancy traits did not show any correlation with first two ordination axes of PCA (Fig. 3, S3). Neither dormancy trait showed any spatial autocorrelation (all Moran’s I correlograms had P > 0.40, not shown).

Separate analyses of relationships between each dormancy trait and each bioclimatic and soil variable showed that only one dormancy trait (PI_PY_) was significantly correlated with more environmental variables while other dormancy traits were either not correlated or showed weak correlations with some environmental variables (Fig. S3, S4, S5). Specifically, PI_PY_ was clearly related to the gradient of aridity, i.e. PI_PY_ increases with increasing temperatures (both mean and maximal summer), decreasing inter-annual variation in mean temperature of warmest quarter, decreasing precipitation and decreasing available soil water capacity (Fig. 3, S3). However, there were two climatic variables, i.e. *IV*BIO5 and *IV*BIO10, which showed significant correlations with majority of dormancy traits (Fig. 3, Table S5). Specifically, final PY dormancy (FPYD, FPYD_25_, FPYD_35_) slightly increased with increasing inter-annual variation in temperatures of warmest quarter (all r = 0.18*–0.19*).

Using cluster analysis, *Medicago* accessions were grouped into four macro-environmental clusters (Fig. 3, 4) clearly differentiated by climatic and soil conditions (Table 3S). Temperature affected PY dormancy (FPYD) depending on clusters (K). Considering each year separately, accessions from aridity conditions (K1 and K4) consistently showed higher FPYD at 25/15°C and lower at 35/15°C. In contrast, accessions from K2 did not change significantly in response to temperature (Fig. 5).

**Figure 4.**
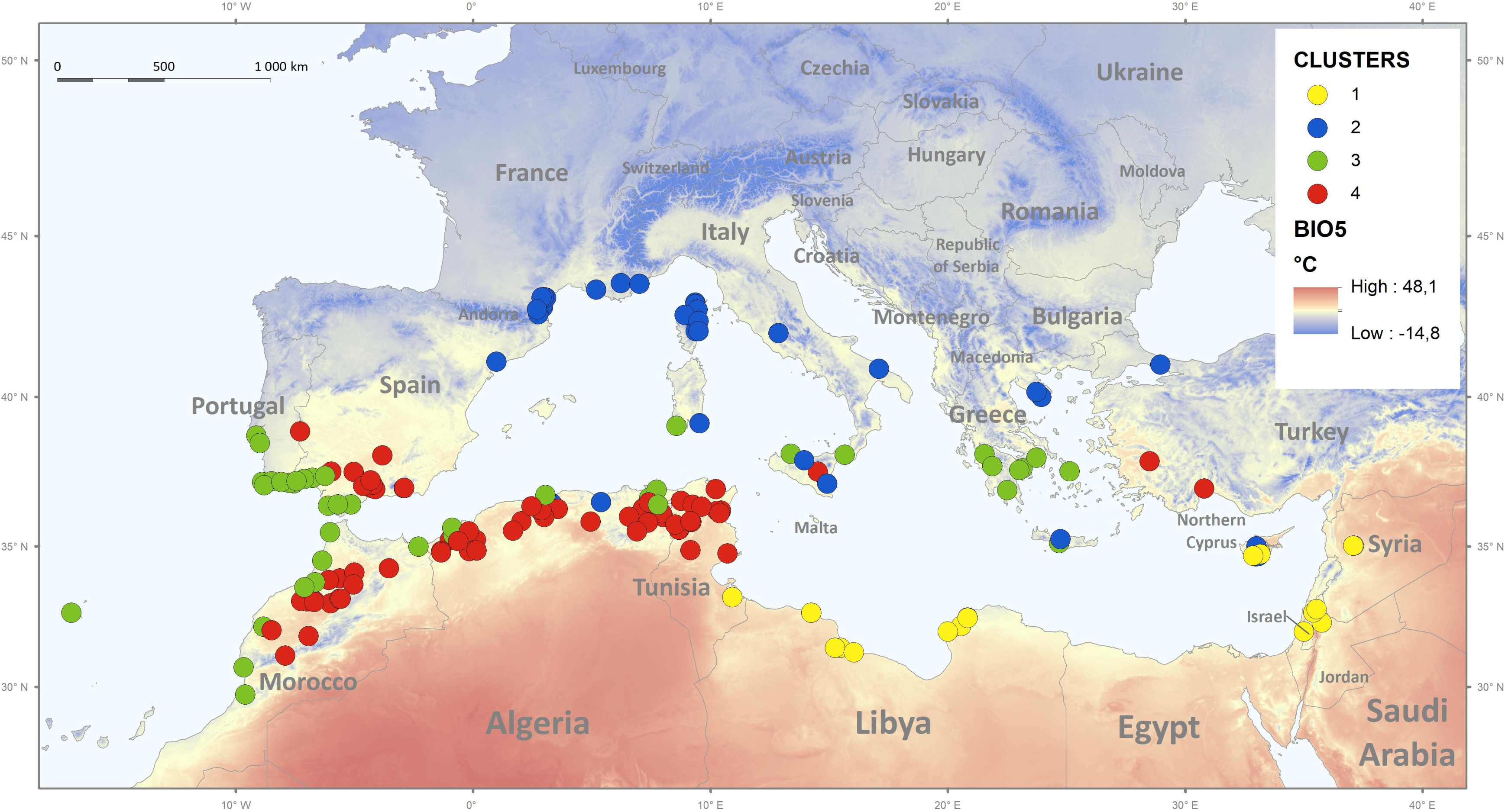
Geographic distribution of studied *M. truncatula* accessions classified in four clusters based on climatic and soil conditions, using Ward’s minimum-variance linkage of Euclidean distance. Grey dots indicate K1, green K2, light blue K3, and yellow K4 cluster, placed on the background of BIO5 (precipitation in the wet quarter).

**Figure 5.**
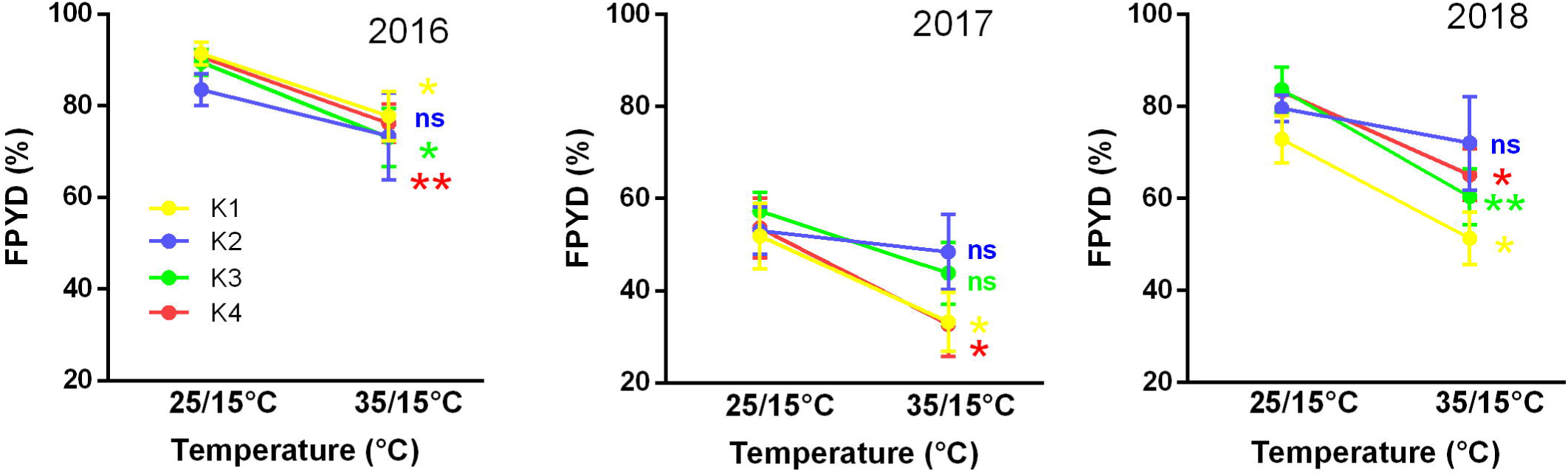
Reaction norms to changes in temperature on final PY dormancy of seeds from K1 to K4 in three years (2016, 2017 and 2018). Vertical bars indicate ± s.e.m. ‘**’ indicates significance at p<0.01, at p<0.05, and ‘nś not significant difference (p > 0.05) between different mean for each cluster respect to temperature treatments.

### Association analysis of dormancy traits

In order to identify molecular mechanisms underlaying physical dormancy and its adaptability, we performed genome-wide association analyses for all dormancy traits (FPYD_25_, FPYD_35_, AUC_25_, AUC_35_, AUC_35-25_, Pl_PY_, Pl_AUC_) and three bioclimatic variables (BIO1, BIO9, BIO12) on 178 accessions. Corresponding Manhattan plots for these analyses are provided in Fig. S6. QQ plots confirmed that FarmCPU was a more suitable model to perform association studies (Fig. S7). Most significant QTNs were identified with AUC_25_, AUC_35-25_, FPYD_25_, Pl_AUC_ and all three bioclimatic variables. To provide a list of significant QTNs, we defined a threshold of 10^-7^ (except for PLA_PY we used a threshold of 10^-4^) (Table S6). 136 candidate genes were identified as potential regulators of physical dormancy (Table S6). A large proportion of candidate genes was annotated as involved in synthesis of secondary metabolites, in cell wall modification and hormone regulation. We performed an over-representation analysis with these 136 candidate genes using a hypergeometric test with Bonferroni correction and we revealed three biological functional GO classes statistically overrepresented (Table S7) and acting as potential regulator of dormancy: response to oxidative stress (GO:0006979), oxidation reduction (GO:0055114), response to chemical stimulus (GO:0042221). Candidate genes belonging to these three GO classes are indicated in Table 1.

**Table 1.** List of QTN identified by GWA studies fo each dormancy trait and belonging to one of three biological function over-represented in our complete list of candidate QTNs. Corresponding chromosome locations and p-values of QTNs are indicated as well as the closest gene ID within +/- 10 kb genomic interval and its corresponding annotations (Mtv5 annotations, Mtv4 annotations, gene ontology and gene description). For complete list and description of all identified QTN see Tables S6 and S7.

| Chrom | Position of QTN | P-value | Candidate gene ID within +/- 10kb interval (Mtv5) | Gene ID v4 | Gene annotation | Gene description |
| --- | --- | --- | --- | --- | --- | --- |
| 1 | 34656783 | 4.25E-11 | MtrunA17Chr1g0184131 | Medtr1g070110 | Hyoscyamine 6-dioxygenase | secondary metabolism.flavonoids.dihydroflavonols |
| 1 | 49585766 | 1.36E-07 | MtrunA17Chr1g0204011 | Medtr1g101830 | Peroxidase | misc.peroxidases |
| 1 | 50761951 | 2.79E-07 | MtrunA17Chr1g0205571 | Medtr1g104590 | Primary-amine oxidase | misc.oxidases - copper, flavone etc |
| 1 | 50761951 | 4.23E-08 | MtrunA17Chr1g0205541 | Medtr1g104550 | Primary-amine oxidase | misc.oxidases - copper, flavone etc |
| 2 | 10895258 | 5.60E-09 | MtrunA17Chr2g0291751 | Medtr2g028980 | Peroxidase | misc.peroxidases |
| 2 | 12184616 | 5.76E-09 | MtrunA17Chr2g0293411 | Medtr2g031920 | Ent-kaurenoic acid oxidase 2 | hormone metabolism.gibberelin.synthesis-degradation.ent-kaurenoic acid hydroxylase/oxygenase |
| 4 | 53502844 | 5.22E-08 | MtrunA17Chr4g0061311 | Medtr4g109360 | UDP-glucose 6-dehydrogenase | cell wall.precursor synthesis.UDP-Glc dehydrogenase (UGD) |
| 4 | 53502844 | 3.62E-08 | MtrunA17Chr4g0061381 | Medtr4g109470 | Flavonoid 3'-monooxygenase | secondary metabolism.flavonoids.dihydroflavonols.flavonoid 3-monooxygenase |
| 4 | 61245281 | 2.56E-07 | MtrunA17Chr4g0072091 | Medtr4g127670 | Peroxidase | misc.peroxidases |
| 5 | 4768980 | 2.23E-12 | MtrunA17Chr5g0400421 | Medtr5g014250 | Beta-amyrin 11-oxidase-like | hormone metabolism.gibberelin.synthesis-degradation.ent-kaurenoic acid hydroxylase/oxygenase |
| 5 | 33045218 | 2.41E-07 | MtrunA17Chr5g0432331 | Medtr5g074710 | Peroxidase | misc.peroxidases |
| 5 | 33045218 | 2.41E-07 | MtrunA17Chr5g0432361 | Medtr5g074770 | Peroxidase | misc.peroxidases |
| 6 | 28332981 | 4.21E-10 | MtrunA17Chr6g0474391 | Medtr6g464620 | Gibberellin 3-beta-dioxygenase | hormone metabolism.gibberelin.synthesis-degradation.GA20 oxidase |
| 6 | 34330903 | 7.87E-06 | MtrunA17Chr6g0479681 | Medtr6g072490 | Cytokinin hydroxylase-like | misc.cytochrome P450 |
| 8 | 6442861 | 1.76E-09 | MtrunA17Chr8g0343001 | Medtr8g018650 | Seed linoleate 9S-lipoxygenase | hormone metabolism.jasmonate.synthesis-degradation.lipoxygenase |
| 8 | 48706047 | 1.32E-09 | MtrunA17Chr8g0391921 | Medtr8g105630 | Glutathione peroxidase | redox.ascorbate and glutathione.glutathione |

## Discussion

*Medicago truncatula* is a representative of species adapted to typical Mediterranean climate characterized by strong seasonality with hot and dry summer often with large diurnal temperature oscillations followed by main rainfall autumn season. In particular, eastern and southern zones of Mediterranean-desert transitions are associated with increased aridity (Thompson, 2005). Summer drought limits growth and is the major cause of seedling mortality, while winter cold limits vegetative growth in certain places. High oscillating temperatures are hypothesized (Baskin and Baskin, 2014) to be one of the main triggers of PY dormancy release and it was confirmed experimentally in several legume species, including *Lupinus, Trifolium, Pisum* and *Medicago* (Hradilová *et al*., 2019; Quinlivan 1961, 1966, 1967, 1968; Taylor, 2005; this study). However, these studies tested only the effect of temperature variation, while in nature more factors vary (Finch-Savage and Footitt, 2017). Especially soil moisture oscillation is very difficult if impossible to mimic in laboratory conditions. The effect of water potential on seed germination was tested only as a static component (Hu *et al*., 2015; Tribouillois *et al*., 2016). As the result, our experimental setup could reveal only temperature related dormancy release.

Variation in germination strategy is particularly relevant for plants inhabiting unpredictable environments and is consistent with seed function, securing next generation in time and space (Finch-Savage and Leubner-Metzger, 2006). Despite being plastic and influenced by range of environmental cues acting both at pre- and post-dispersal, the seed dormancy including of physical type (Dias *et al*., 2011; Hudson *et al*., 2015; Hradilová *et al*., 2017) is highly heritable trait. In our study, seed dormancy release varied among genotypes and years and this could potentially act as a mechanism that favours the persistence of the seed in the soil and helps to distribute genetic diversity through time (Hudson *et al*., 2015; Long *et al*., 2015).

### Association between seed dormancy traits and the environment

In this work we have tested if seed dormancy release in *M. truncatula* contributes to local adaptation to diverse environmental conditions across the species geographic range. However, when taking average estimates of various traits characterizing absolute dormancy release (i.e., FPYD, AUC and AUC_35-25_), no relationships were found between climatic or environmental conditions and mean dormancy status per accession (Figure 3). Our observations are in agreement with study of Mediterranean wild lupines (Berger *et al*., 2017) or perennial woody legume (*Vachellia aroma*) along a precipitation gradient (Ferreras *et al*., 2017) that did not find any relationship between climate and dormancy release. In contrast, *A. thaliana* (Debieu *et al*., 2013; Postma and Agren, 2015) and other winter-annual species such as *Betta vulgaris* subsp*. maritima*, *Biscutella didyma*, *Bromus fasciculatus* and *Pisum sativum* subsp. *elatius* showed a cline in dormancy (Ferreras *et al*., 2017; Hradilová *et al*., 2019; Lampei *et al*., 2017; Wagmann *et al*., 2012). In particular, more dormant genotypes of mentioned taxa occurred in lower latitudes or more arid habitats with seasonally unpredictable precipitation and less-dormant in higher latitudes or more humid habitats, suggesting dormancy as adaptation securing populations survival in less predictable conditions. However, in arid and unpredictable environments, there are also species called ‘risk-takin’ with lower dormancy and rapid germinate in response to lower rainfall events. The high and reliable seed production determines that the consequences of failure to establish in these species are less dire (Duncan *et al*., 2009).

Does the absence of relationship between average estimates of dormancy release in *Medicago* and observed climatic clines within dataset suggest that selection do not operate on dormancy traits? Taking into account plasticity of dormancy release between temperature treatments and among years, however, revealed that physical dormancy plasticity (PI_PY_) increases with increasing aridity (Fig. 3, S3), In addition, the accessions grouped by environmental clusters helped to explain the interpretations based on the reaction norm, where the accessions from aridity conditions (K1 and K4) consistently response to temperature (Fig. 5). It follows, that more plastic behaviour can distribute germination across the year and acts as a bet-hedging strategy (Burghardt *et al*., 2016), suggesting that under more unpredictable environmental conditions genetic components of phenotypic variance may be lower and thus a reduced evolutionary response to selection would be possible (Hudson *et al*., 2015). The bet-hedging strategy is thus positively associated with more arid habitat as found in pea (Hradilová *et al*., 2019) and plastic responses provide potential to cope with high levels of environmental heterogeneity (Cochrane, 2019).

We have shown that inter-annual summer temperatures relate to the absolute dormancy values. More unstable summer temperatures between years tend to increase the PY dormancy levels (Table 5S) and might contributes to avoid a high percentage of germination under adverse temperatures condition or the “false breaks” (seed germination outside the optimal growing season) (Taylor, 2005).

### Potential shortcomings of the study

Although the dormancy is genetically determined, it also depends on the environmental conditions experienced by the mother plant and the subsequent status of the seed (Finch-Savage and Footitt, 2017; Hudson *et al*., 2015). This was shown in several taxa, including *Trifolium* and *Medicago* (Collins, 1981; Donnelly *et al*., 1972; Hudson *et al*., 2015; Jaganathan, 2016; Quinlivan, 1965, 1966; Quinlivan and Millington, 1962; Taylor, 1996; Tozer and Ooi, 2014). There are several levels of possible sources of variability, from effect of maternal plant status (drought, photoperiod, nutrition) to variability between individuals from the same population or even the same individual (Penfield and MacGregor, 2017). Distinguishing between maternal and environmental effects is difficult. The information on the maternal environmental can be mediated via the nutrition, phytohormones or at gene expression levels and variability in the seed sensitivity (Finch-Savage and Footitt, 2017). All this might contribute to *M. truncatula* seed stock generated in our study. To minimize environmental maternal effects, we grew the accessions under common garden conditions (glasshouse), but these were to some extend variable between years. In 2016, the accessions were sown February to April and harvested in hot July with day temperature over 35°C, while in 2017 and 2018 there were sown in September and grew over winter, flowered February and maturate in April, with day temperatures in glasshouse around 28°C. The higher temperature in 2016 during the seed filling period resulted in more dormant seeds in some accessions in relation to 2017 and 2018 (Fig. 6). In addition to this, seeds from different accessions differ up to 3 weeks in maturation due to differences in flowering time and also the individual seed stock from given accession was harvested in period of about 3 weeks. This need to be considered in follow up studies. We can also just speculate what is the influence of environmental conditions at the original site of the given accessions. This could be tested by either *in situ* collection of the seeds or reciprocal common garden experiments.

**Figure 6.**
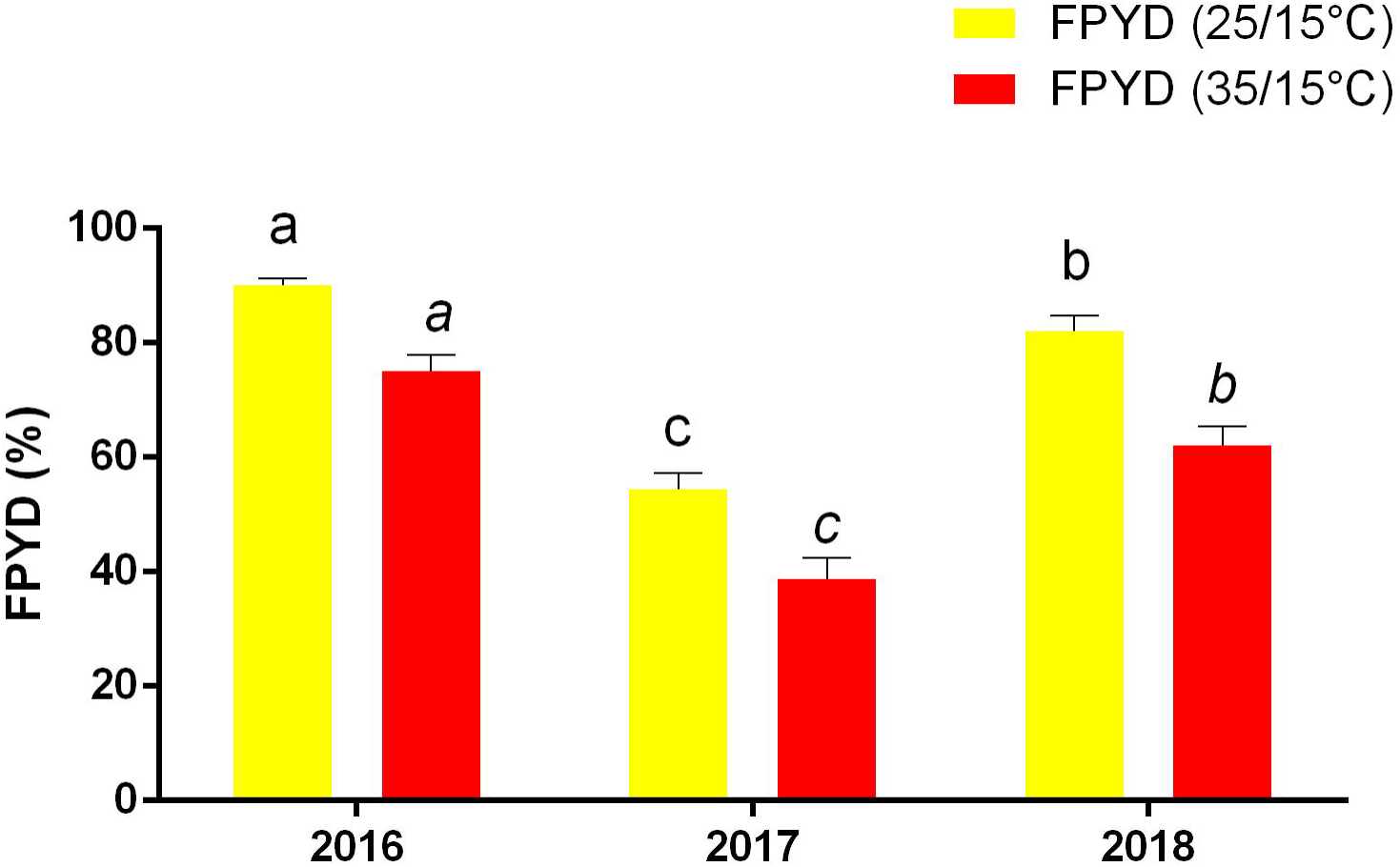
Final PY dormancy (FPYD) of seeds in *Medicago truncatula* accessions during 2016, 2017 and 2018 at 25/15°C and 35/15°C. Letters were used to indicate the differences between years for each temperature.

Our analysis was also likely impacted by several factors inherent to available *Medicago* set. At first, there is substantial imprecision in the GPS localization of the origin of some accessions (Stine and Hunsaker, 2001) leading consequently to incorrect extracted environmental factors (Brus *et al*., 2018; Buisson *et al*., 2010; Meyer *et al*., 2016). Secondly, there is geographical bias towards the western part of Mediterranean with underrepresented parts of the native species range, such as Italy, Adriatic Sea coast, Turkey, Lebanon and Israel. In addition the character of WorldClim data (average in term of time and also space) masks the micro-ecological pattern and in geographically complex regions, environmental conditions change considerably over short spatial scales, such that neighbouring populations can be subject to different selective pressures as found in the study of seed dormancy of Swedish *A. thaliana* accessions (Kerdaffrec and Nordborg, 2017). We compared the genetic groups identified by Gentzbittel *et al*. (2019) to our study macroclimatic clusters and found significant correspondence (*r^2^*=0.582**, Table S5). As discussed in Gentzbittel *et al*. (2019), geography explains 5% of variation, while climate is the major source of variation (41.5%).

### Genetic basis of seed dormancy release in legumes

In contrast to the seed development, the genetic basis of *Medicago* seed germination was studied only by Dias *et al*. (2011). These authors however focused on true germination, e.g. radicle emergence, removing the seed coat prior to testing, assessing thus physiological dormancy, while we were interested in PY dormancy executed by seed coat permeability. Since the dormancy is indirectly evaluated as the release from dormancy by imbibition and germination, the detected candidate genes might be related to the changes in seed coat mediated imbibition process rather than dormancy per se. There was no overlap in associated loci between the tested seed dormancy traits, despite the detection of similar set of candidate genes (Table S8). This is similar to other tested quantitative and complex traits such as drought and biomass (Kang *et al*. 2015) where different traits had different candidate genes.

We have detected four genes active in flavonoid and phenylpropanoid biosynthesis pathway leading either to flavonoids or via polymerization to lignins (Francoz *et al*., 2018; Smýkal *et al*., 2014) impregnating the seed coat. Notably, homologues genes were detected when compared dormant and non-dormant pea seed coat expression (Hradilová *et al*., 2015). Furthermore, hydrolytic enzymes such as xyloglucan 6-xylosyltransferase, xylogalacturonan beta-1,3-xylosyltransferase involved in plant cell wall modification were identified. Notably, one of the QTL identified in biparental mapping of *Medicago* seed germination also encodes xyloglucan endotransglucosylase (Dias *et al*., 2011). The β-1,3-Glucanase (EC 3.2.1.39) plays roles in the regulation of seed germination, dormancy and in the defence against pathogens (Leubner-Metzger, 2003). β-1,3-glucan layer is in seed coat of cucurbitaceous species and confers seed semipermeability (Leubner-Metzger, 2003). In tobacco seeds, β-1,3-Glucanase was shown to be at micropylar part of the endosperm prior to radicle protrusion, and seems to act into cell wall loosening (Leubner-Metzger *et al*., 1995).

Pectinesterases were isolated from germinating seeds of various species and are assumed to play an important role in loosening cell walls (Nighojkar *et al*., 1994).

Polygalacturonases (EC 3.2.1.15) are another cell wall degrading enzymes. These were shown to play an essential role in pollen maturation and in pectin metabolism during fruits softening and weakening of the endosperm cell walls (Sitrit *et al*., 1999). Exo-(1→4)-β-galactanases (EC 3.2.1.23) play various roles in physiological events, including cell wall expansion and degradation during soft fruit ripening and were found to be involved in the mobilization of polysaccharides from the cotyledon cell walls of *Lupinus angustifolius* following germination (Buckeridge *et al*., 2005).

Nitric oxid (NO) was recently shown to be involved in plant development including seed germination (Hancock and Neils, 2019). NO-dependent protein post-translational modifications are proposed as a key mechanism underlying NO signalling during early seed germination. Our GWAS analysis identified seven putative peroxidase and thio-/peroxiredoxin genes. Peroxiredoxins (EC 1.11.1.15) catalyze the reduction of hydroperoxides, conferring resistance to oxidative stress. Recent studies have demonstrated that ROS have key roles in the release of seed dormancy, as well as in protection from pathogens (Müller *et al*., 2009; Jeevan Kumar *et al*., 2015; Raviv *et al*., 2018). Thioredoxins were identified to promote seed germination (in Bewley *et al*., 2013). Peroxidases (EC 1.11.1.7) are also implicated in lignin/suberin formation during the polymerization of monolignols (Novo-Uzal *et al*., 2013) synthetized in of the final steps of the phenylpropanoid pathway. Phytohormones and especially gibberellins are known to play important roles in seed development and germination (reviewed in Graebner *et al*., 2012), and since *Medicago* seeds have both physical and physiological dormancy, it is not surprising to find gibberellin 20-oxidase and two ent-kaurenoic acid oxidase genes to be associated with dormancy release (AUC_25) or environmental factors (BIO12), respectively. Genomic signature of *M. truncatula* adaptation to climate was studied by Yoder *et al*. (2014) using essentially same set of lines. They analysed relationship to BIO1, BIO3 and BIO16 so there is the overlap in only BIO1 (annual mean temperature) with our study. The different candidate genes were detected. This could be attributed to differences in accessions, analytical methods as well as *Medicag*o genome versions. However despite these differences, some similar genes were detected such as 1,3-glucanase or kinases. Several kinases and disease resistance (TIR-NBS-LRR class) genes associated with dormancy release traits. These have been implicated in pathogen sensing and host resistance, which might reflect the sensing of cell wall modulating enzymes activities, similar to pathogen attack (Pollard, 2018). As the result of bet-hedging, seeds in the soil form long-term seed bank where they need to be protected from microbial decay by presence of secondary metabolites as well as seed defence enzymes (Pollard, 2018; Raviv *et al*., 2018). Therefore, one of the possible future directions of seed dormancy release studies should go into study of seed-soil-microbiome relationships and seed coat enzymatic activities.

## Supplementary data

Fig. S1. Geographic distribution of studied *M. truncatula* accessions. Fig. S2. Frequency distributions of dormancy traits.

Fig. S3. Correlation between the phenotypic plasticity index of final dormancy (PI_PY_) and selected environmental variables.

Fig. S4. Correlation chart of the dormancy release traits and environmental variables.

Fig. S5. Correlation chart of the dormancy release traits, soil variables and inter-annual climatic variables.

Fig. S6. Manhattan plots of mapped SNP markers associated with dormancy or bioclimatic traits.

Fig.S7. Quantile–quantile (Q-Q) plots for all the traits obtained by standard mixed linear model (EMMA) and multi-locus linear model (FarmCPU).

Table S1. List of tested *Medicago* accessions with calculated seed dormancy traits and extracted environmental variables.

Table S2. Basic descriptive statistics of 23 bioclimatic variables and 10 soil variables of sites of accessions origin.

Table S3. Classification of 176 *M. truncatula* accessions in four cluster based on environmental and climatic conditions.

Table S4. Pearson coefficients-probabilities between dormancy traits and bioclimatic variables.

Table S5. Regression coefficient (r^2^) between environmental variables and plasticity index by macroecological and genetic clusters of Gentzbittel *et al*. (2019).

Table S6. Complete list of QTN identified by GWA studies for each dormancy trait.

Table S7. Over-representation analysis of the 136 candidate genes potentially involved in dormancy traits.

## Acknowledgements

This work was funded by the Grant Agency of the Czech Republic, 16-21053S.

